# Cell entry-independent role for the reovirus μ1 protein in regulating necroptosis and the accumulation of viral gene products

**DOI:** 10.1101/541623

**Authors:** Katherine E Roebke, Pranav Danthi

## Abstract

The reovirus outer capsid protein μ1 regulates cell death in infected cells. To distinguish between the role of incoming, capsid-associated and newly synthesized μ1, we used siRNA-mediated knockdown. Loss of newly synthesized μ1 protein does not impact apoptotic cell death in HeLa cells but enhances necroptosis in L929 cells. Knockdown of μ1 also impacts aspects of viral replication. We found that while μ1 knockdown results in diminished release of infectious viral progeny from infected cells, viral minus strand RNA, plus strand RNA, and proteins that are not targeted by the μ1 siRNA accumulate to a greater extent when compared to control siRNA-treated cells. Furthermore, we observe a decrease in sensitivity of these viral products to inhibition by GuHCl (which targets minus strand synthesis to produce dsRNA) when μ1 is knocked down. Following μ1 knockdown, cell death is also less sensitive to treatment with GuHCl. Our studies suggest that the absence of μ1 allows enhanced transcriptional activity of newly synthesized cores and the consequent accumulation of viral gene products. We speculate that enhanced accumulation and detection of these gene products due to a μ1 knockdown potentiates RIP3 dependent cell death.

**IMPORTANCE:** We use mammalian reovirus as a model to study how virus infections result in cell death. Here, we sought to determine how viral factors regulate cell death. Our work highlights a previously unknown role for reovirus outer capsid protein μ1 in limiting the induction of a necrotic form of cell death called necroptosis. Induction of cell death by necroptosis requires the detection of viral gene products late in infection. μ1 limits cell death by this mechanism because it prevents excessive accumulation of viral gene products that trigger cell death.

## INTRODUCTION

Replication of a virus within host cells can result in cell death (1), which is often detrimental to virus infection (2). To counter the antiviral effect of cell death, many viruses encode proteins which prevent cell death (1, 2). In some cases, cell death can have pathogenic consequences. In instances where cell death is required for cell-to-cell spread of the virus or dissemination of virus beyond the primary site of replication, cell death can exacerbate disease (1). Additionally, tissue injury caused by cell death can also contribute to viral disease. Mammalian reovirus has been used as an experimental model to investigate the impact of cell death on viral disease. In a mouse model of infection, the potential for reovirus to cause heart or CNS disease is dependent on its capacity to induce cell death by apoptosis (3-5). Numerous studies have mapped out both viral and cellular factors that influence apoptosis induction and disease following reovirus infection.

The known viral determinants of reovirus-induced apoptosis are the attachment protein σ1 and the membrane penetration protein μ1 (6-9). The role of σ1 in controlling cell death correlates with the efficiency with which σ1 mediates attachment to host cell receptors (8, 9). The reovirus outer-capsid protein μ1, which plays an essential role in delivering the viral core particle into the cytoplasm, is also important for controlling the efficiency with which reovirus induces apoptosis. Mutations within the C-terminal region of μ1 diminish the capacity of reovirus to evoke proapoptotic signaling (10, 11). Cell death signaling following ectopic expression of μ1 in plasmid transfected cells also supports this idea (12, 13). Even in this artificial context, the C-terminal region of μ1, which affects the localization of μ1 to intracellular membranous structures such as mitochondria, plays an important role in release of proapoptotic factors cytochrome c and smac/DIABLO from the mitochondria and activation of effector caspases (12, 13). Based on the evidence that events that occur prior to viral gene expression are sufficient for the induction of apoptosis (9, 14), it is assumed that the effect of μ1 on the apoptotic potential of reovirus is related to the function of μ1 as part of incoming viral capsid. However, this idea has not directly been tested.

Depending on the cell type, reovirus can elicit another form of regulated cell death called necroptosis (15, 16). Unlike apoptosis, necroptosis is thought to be an inflammatory form of cell death (17). Reovirus-induced necroptosis is initiated by sensing of incoming genomic dsRNA by RIG-I-like-receptors (RLRs) retinoic acid-inducible gene I (RIG-I) and melanoma differentiation-associated protein 5 (MDA5) (16). RLRs signal via the mitochondrial antiviral signaling protein (MAVS), to produce type 1 interferon (IFN). In addition to IFN signaling, *de novo* synthesis of viral dsRNA genome is required for induction of necroptosis (16, 18). Together, these events in reovirus infection lead to receptor interacting protein 1 (RIP1) and RIP3 dependent cell death (15, 16). The necroptotic effector protein mixed linkage kinase like protein (MLKL) is also activated at times that are consistent with the induction of cell death (16). Our working hypothesis is that *de novo* synthesized genomic RNA or its products are detected by an IFN-stimulated gene (ISG) to induce necroptosis. Viral factors that increase dsRNA synthesis or control the exposure of dsRNA are likely to influence necroptosis. However, no such factors have yet been identified.

We sought to identify viral factors that contribute to the induction of cell death following reovirus infection. Given the previously described role of μ1 in cell death, we aimed to further dissect the mechanisms by which μ1 is involved in reovirus-induced cell death. Here, we explored the role of newly synthesized μ1 in cell death by using siRNA mediated knockdown. We observed that knockdown of μ1 does not impact apoptosis induction by reovirus suggesting that μ1 present in the incoming capsid is sufficient to regulate apoptosis. In contrast, knockdown of μ1 accelerates necroptosis induction following reovirus infection, indicating that newly synthesized μ1 impacts this form of cell death. Furthermore, we discovered that knockdown of the μ1 protein in infected cells results in an increased accumulation of progeny dsRNA, secondary transcripts produced from dsRNA, and viral proteins in the infected cells. These data highlight a new function for newly synthesized μ1 in controlling the levels of viral gene products in infected cells and support the model that viral components that are synthesized late in infection are detected to elicit necroptotic cell death.

## MATERIALS AND METHODS

### Cells

L929 cells obtained from ATCC were maintained in Eagle’s MEM (Lonza) supplemented to contain 5% fetal bovine serum (FBS) (Life Technologies), and 2 mM L-glutamine (Invitrogen). Spinner-adapted Murine L929 cells were maintained in Joklik’s MEM (Lonza) supplemented to contain 5% fetal bovine serum (FBS) (Sigma-Aldrich), 2 mM L-glutamine (Invitrogen), 100 U/ml penicillin (Invitrogen), 100 μg/ml streptomycin (Invitrogen), and 25 ng/ml amphotericin B (Sigma-Aldrich). Spinner-adapted L929 cells were used for cultivating, purifying and titering viruses. HeLa cells obtained from Melanie Marketon’s lab at Indiana University were maintained in Dulbecco’s MEM (Lonza) supplemented to 5% FBS (Sigma-Aldrich, and 2mM L-glutamine (Invitrogen).

### Reagents

Q-VD-OPh was purchased from Cayman Chemical Company. GuHCl and Ribavirin were purchased from Sigma-Aldrich. Custom synthesized siRNAs were purchased from Dharmacon. siRNA targeting β-galactosidase was used as control. The siRNA sequences used are as follows: β-galactosidase – CUACACAAAUCAGCGAUUU, μ1 – GGAAAGAGUUAUAAAGAGA, and σ3 – GCGCAAGAGGGAUGGGACA, and Antisera raised against reovirus capsid and σNS were obtained from T. Dermody and have been previously described (19). Monoclonal antibody against IFNAR and rabbit antisera against RIP3 were purchased from Santa Cruz Biotechnology and Pro-science respectively. Mouse antiserum specific for PSTAIR was purchased from Sigma. Alexa Fluor-conjugated anti-mouse IgG and anti-rabbit IgG secondary antibodies were purchased from Invitrogen. IR-conjugated anti-guinea pig IgG was purchased from LICOR.

### Virus purification

Prototype reovirus strains of T3D, T1L, and T1L/T3DM2 were regenerated by plasmid based reverse genetics <u>(20)</u>. Purified reovirus virions were generated using second- or third-passage L-cell lysates stocks of reovirus. Viral particles were Vertrel-XF (Dupont) extracted from infected cell lysates, layered onto 1.2- to 1.4-g/cm^3^ CsCl gradients, and centrifuged at 187,183 x *g* for 4 h. Bands corresponding to virions (1.36 g/cm^3^) were collected and dialyzed in virion-storage buffer (150 mM NaCl, 15 mM MgCl_2_, 10 mM Tris-HCl [pH 7.4]) <u>(21)</u>.

### siRNA transfection

In 96-well plates, 0.25 μl Lipofectamine 2000 was used to transfect siRNA to a final concentration of 100 nM, or 0.75µl INTERFERin (Polyplus) was used to transfect siRNA to a final concentration of 5 nM. Cells (1 × 10^4^) were seeded on top of the siRNA-lipofectamine/INTERFERin mixture. In 24-well plates, 1.5 µl Lipofectamine 2000 was used to transfect siRNA to a final concentration of 42 nM or 3 µl of INTERFERin was used to transfect siRNA to a final concentration of 15 nM. Cells (1 x 10^5^) were seeded on top of the siRNA-lipofectamine/INTERFERin mixture. Virus infection was performed 28-48 h following siRNA transfection.

### Infection and preparation of extracts

Cells were either adsorbed with PBS or T3D at room temperature for 1 h, followed by incubation with media at 37°C for the indicated time interval. When needed, DMSO, Ribavirin, GuHCl, Z-VAD-FMK, Q-VD-OPh, or anti-IFNAR Ab were added to the media immediately after the 1 h adsorption period. For preparation of whole cell lysates, cells were washed in phosphate-buffered saline (PBS) and lysed with 1X RIPA (50 mM Tris [pH 7.5], 50 mM NaCl, 1% TX-100, 1% DOC, 0.1% SDS, and 1 mM EDTA) containing a protease inhibitor cocktail (Roche) and 2 mM PMSF, followed by centrifugation at 15000 × *g* for 10 min to remove debris.

### Immunoblotting

Cell lysates were resolved by electrophoresis on 10% polyacrylamide gels and transferred to nitrocellulose membranes. Membranes were blocked for at least 1 h in TBS containing 5% milk, or T20 Starting Block (Thermo Fisher) and incubated with antisera against σNS (1:2000), reovirus (1:10000), RIP3 (1:1000), or PSTAIR (1:10000) at 4°C overnight. Membranes were washed three times for 5 min each with washing buffer (TBS containing 0.1% Tween-20) and incubated with 1:20000 dilution of Alexa Fluor conjugated goat anti-rabbit IgG (for reovirus polyclonal and RIP3), goat anti-mouse IgG (for PSTAIR), or IRDye-conjugated anti-guinea pig IgG (for σNS) in blocking buffer. Following three washes, membranes were scanned using an Odyssey Infrared Imager (LI-COR) or Chemidoc MP Imaging System (Biorad). Band intensity was analyzed using Image Studio Lite software (LICOR).

### Quantitation of cell death by Acridine orange-ethidium bromide (AOEB) staining

ATCC L929 cells (2×10^5^) grown in 96 well-plates were adsorbed with the indicated MOI of reovirus at room temperature for 1 h. The percentage of dead cells at the indicated time following infection was determined using AOEB staining as described previously (6). For each experiment, >200 cells were counted, and the percentage of isolated cells exhibiting orange staining (EB positivity) was determined by epi-illumination fluorescence microscopy using a fluorescein filter set on an Olympus IX71 microscope.

### Assessment of viral growth

L929 cells in 96-well plates were adsorbed in triplicate infections with 2 PFU/cell of T3D for 1 h. Cells were washed once with PBS and incubated in supplemented EMEM at 37°C for 24 h. Twice, the cells were frozen at −80°C and then thawed prior to determination of viral titer by plaque assay. Viral yields were calculated according to the following formula: log_10_yield_24h_ = log_10_(PFU/ml)_24h_-log_10_(PFU/ml)_0h_.

### Plaque assays

Plaque assays to determine infectivity were performed as previously described with some modifications (21, 22). Briefly, control or heat-treated virus samples were diluted into PBS supplemented with 2 mM MgCl_2_(PBS^Mg^). L cell monolayers in 6-well plates (Greiner Bio-One) were infected with 100 μl of diluted virus for 1 h at room temperature. Following the viral attachment incubation, the monolayers were overlaid with 4 ml of serum-free medium 199 (Sigma-Aldrich) supplemented with 1% Bacto Agar (BD Biosciences), 10 μg/ml TLCK-treated chymotrypsin (Worthington, Biochemical), 2 mM L-glutamine (Invitrogen), 100 U/ml penicillin (Invitrogen), 100 μg/ml streptomycin (Invitrogen), and 25 ng/ml amphotericin B (Sigma-Aldrich). The infected cells were incubated at 37°C, and plaques were counted 5 d post infection.

### Assessment of caspase-3/7 activity

ATCC L929 cells (1 × 10^4^) were siRNA transfected as described above and seeded into black clear-bottom 96-well plates. 24 h following transfection, cells were adsorbed with 10 PFU/cell of reovirus in serum-free medium at room temperature for 1 h. Following incubation of cells at 37°C for 48 h, caspase-3/7 activity was quantified using the Caspase-Glo-3/7 assay system (Promega). Luminescence was quantified using plate reader Synergy H1 Hybrid Reader (BioTek).

### RT-qPCR

RNA was extracted from infected cells at various time intervals after infection using Total RNA mini kit (Biorad). For RT-qPCR, 0.5 to 2 µg of RNA was reverse transcribed with the High Capacity cDNA Reverse Transcription Kit (Applied Biosystems) using random hexamers for amplification of cellular genes or gene specific primers for amplification of either minus or plus strand viral RNA. Gene specific primers were used at a final concentration of 0.1 µM. A 1:10 dilution of the cDNA was subjected to PCR using SYBR Select Master Mix (Applied Biosystems). Fold increase in gene expression with respect to control sample (indicated in each figure legend) was measured using the ΔΔC_T_method (23). Calculations for determining ΔΔC_T_values and relative levels of gene expression were performed as follows: (i) fold increase in cellular gene expression = 2 ^-[(gene of interest CT – GAPDH CT)24 h – (gene of interest CT – GAPDH CT)0 h]^; (ii) fold increase in viral gene expression = 2 ^-[(T3D S1 CT – GAPDH CT)μ1 siRNA – (T3D S1 CT – GAPDH CT)control siRNA]^; (iii) ratio of plus strand RNA to minus strand RNA = 2 ^-[(T3D S1 CT – GAPDH CT)plus strand RNA – (T3D S1 CT – GAPDH CT)minus stand RNA]^; (iv) Relative levels of RNA following treatment with GuHCl = 2^-[(T3D S1 CT – GAPDH CT)15-50 mM GuHCl – (T3D S1 CT – GAPDH CT)0 mM GuHCl])^.

### Statistical analysis

Statistical significance between experimental groups was determined using the unpaired student’s *t*-test function in excel and graphed using Graphpad Prism software. Statistical analyses for differences in gene expression by RT-qPCR were done on the ΔC_T_ values.

## RESULTS

### Newly synthesized μ1 does not impact reovirus-induced apoptosis

The reovirus outer capsid protein μ1 regulates apoptotic cell death following infection (9-13, 24). However, whether this is a function of incoming capsid-associated μ1 or newly synthesized μ1 protein has not directly been evaluated. To determine if newly synthesized μ1 is responsible for this described role in apoptosis, we knocked down the levels of the T3D μ1 protein in reovirus infected HeLa cells using siRNA. As expected, the siRNA significantly diminished levels of newly synthesized μ1 in infected cells (Figure 1A). To assess the impact of μ1 knockdown on levels of cell death, HeLa cells transfected with either control or μ1 siRNA were infected with T3D for 48 h and cell death was quantified by acridine orange ethidium bromide (AOEB) staining (Figure 1B). We observed that preventing the expression of newly synthesized μ1 did not alter levels of cell death induced in HeLa cells following T3D infection. HeLa cells display markers of apoptosis following reovirus infection and have previously been suggested to undergo cell death by apoptosis following reovirus infection (8, 14, 25). To confirm that cell death in HeLa cells used for these experiments occurred by apoptosis, the infected cells were treated with pan-caspase inhibitor Q-VD-OPh (QVD) (Figure 1C). Cell death in the presence control and μ1 siRNA was significantly decreased by treatment with QVD (Figure 1C). Thus, newly synthesized μ1 does not affect the efficiency of apoptotic cell death following reovirus infection. These data also suggest that the previously suggested role for μ1 in apoptosis is a function of μ1 that is a component of the entering virus particles.

**Figure 1.**
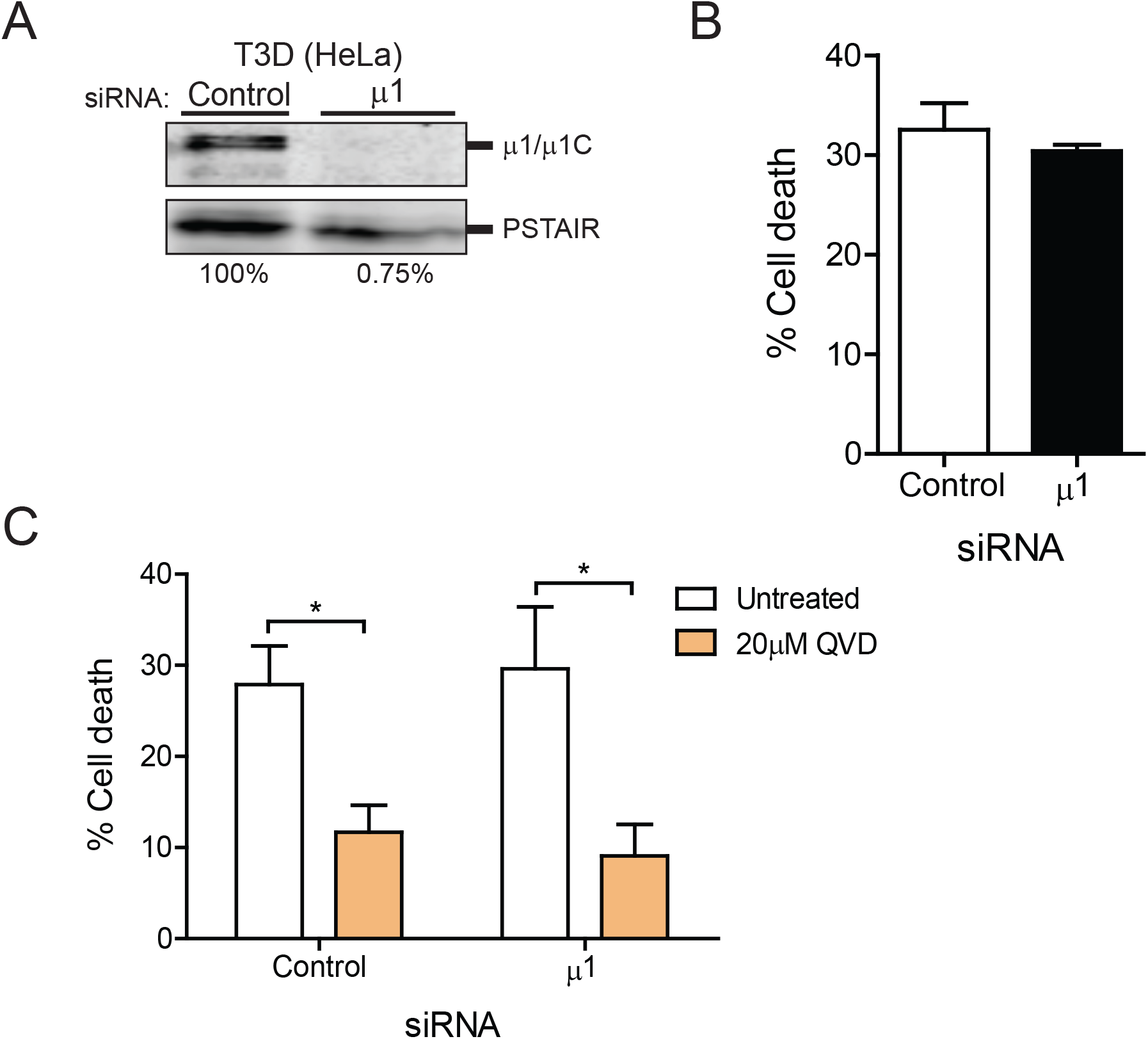
Newly synthesized μ1 does not impact apoptotic cell death. HeLa cells were transfected with either control β-gal or μ1 siRNA using INTERFERin. 24 h following transfection, the cells were infected and processed as described. (A) Cells were infected with T3D at an MOI of 100 PFU/cell. Cell lysates prepared 24 h following infection were immunoblotted for the μ1 protein using anti-reovirus antisera and anti-PSTAIR mAb. Levels of μ1 and μ1C bands relative to PSTAIR are indicated. The level of μ1 and μ1C relative to PSTAIR in control siRNA treated cells was considered 100%. (B) Cells were infected with T3D at an MOI of 100 PFU/cell. At 48 h post infection, cell death was quantified by AOEB staining. (C) Cells were infected with T3D at an MOI of 100 PFU/cell and either left untreated or were treated with 20 µM Q-VD-OPh when infection was initiated. At 48 h post infection, cell death was quantified by AOEB staining. Mean values for three independent infections are shown. Error bars indicate SD. P values were determined by student’s t-test. *, p < 0.05

### Knockdown of newly synthesized μ1 drives an increase in necroptotic cell death

L929 cells succumb to reovirus infection by undergoing a different form of cell death, called necroptosis (16). The requirements for induction of this necrotic form of cell death are distinct from those described for reovirus induced apoptosis (18). To determine if newly synthesized μ1 plays a role in the regulation of necroptotic cell death, we used siRNA-mediated knockdown of μ1 following T3D infection in L929 cells. Analogous to HeLa cells, the siRNA significantly diminished levels of newly synthesized μ1 in infected L929 cells (Figure 2A). To determine if knockdown of μ1 impacts levels of cell death, L929 cells transfected with control or μ1 siRNA were infected with T3D and cell death was quantified by AOEB staining (Figure 2B). Interestingly, in contrast to what we observed in HeLa cells, loss of μ1 in L929 cells enhanced cell death. Even at a time interval where control siRNA treated cells exhibited a minimal level of cell death (~5%) following reovirus infection, μ1 knockdown led to significantly more cell death (~ 30%). These data suggest that newly synthesized μ1 negatively regulates L929 cell death following reovirus infection.

**Figure 2.**
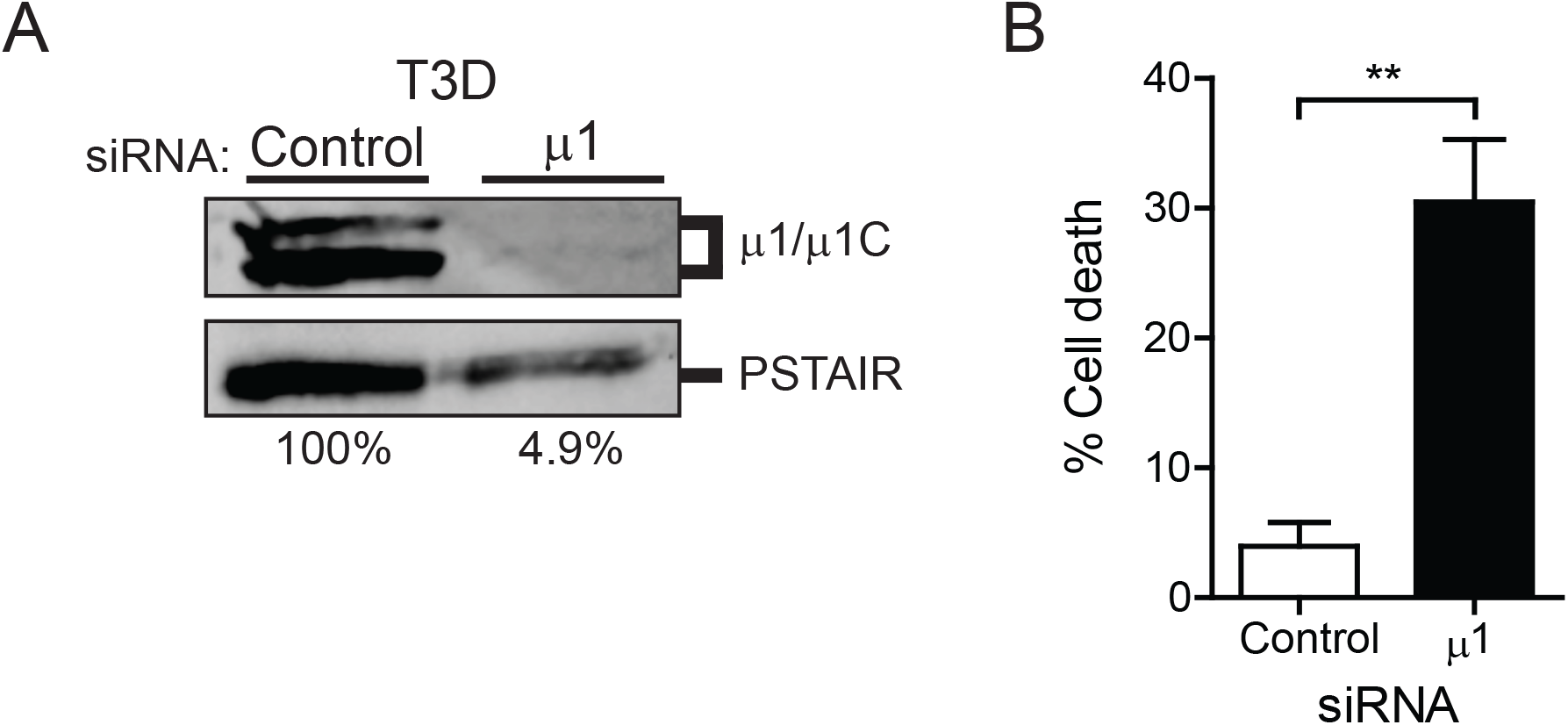
Knockdown of newly synthesized μ1 increases cell death in L929 cells. L929 cells were transfected with either control β-gal or μ1 siRNA using lipofectamine. 24 h following transfection, the cells were infected with T3D at an MOI of 10 PFU/cell and processed as described. (A) Cell lysates prepared 24 h following infection were immunoblotted for the μ1 protein using anti-reovirus antisera and anti-PSTAIR mAb. The level of μ1 and μ1C relative to PSTAIR in control siRNA treated cells was considered 100%. (B) Cell death was quantified 24 h following infection by AOEB staining. Mean values for three independent infections are shown. Error bars indicate SD. P values were determined by student’s t-test. *, p < 0.05; **, p < 0.005

To ensure that the effect of the μ1 siRNA was specific and related to its complementarity to the T3D M2 gene (Figure 3A), we evaluated the capacity of the μ1 siRNA to affect levels of μ1 from prototype strain T1L or a M2 gene reassortant T1L/T3DM2 which expresses T3D μ1 in a T1L genetic background (Figure 3B). We observed that T3D M2 specific siRNA failed to affect expression of T1L μ1 (due to sequence mismatch) but reduced μ1 expression after infection with T1L/T3DM2. Thus, the μ1 siRNA we have designed specifically and efficiently diminishes the level of T3D μ1 protein in infected cells. To confirm that the increased cell death phenotype is specific to knocking down μ1 from T3D, we measured cell death in μ1 siRNA-treated cells infected with T1L or T1L/T3DM2 (Figure 3C). Because T1L induces cell death less efficiently (6, 7, 15), T1L induced cell death was measured at a later time post infection. Regardless, μ1 siRNA failed to affect cell death following T1L infection. In contrast, knockdown of μ1 following infection with T1L/T3DM2 demonstrated an increase in cell death similar to what was observed with T3D. This suggests that the increase in cell death observed is due to the specific siRNA-knockdown of μ1 and not due to off target effects of μ1 siRNA on a cellular signaling pathway. Having established the specificity of the siRNA and the enhanced cell death phenotype observed, we performed the remainder of the experiments with T3D, which has been predominantly used to define reovirus-driven cell death pathways.

**Figure 3.**
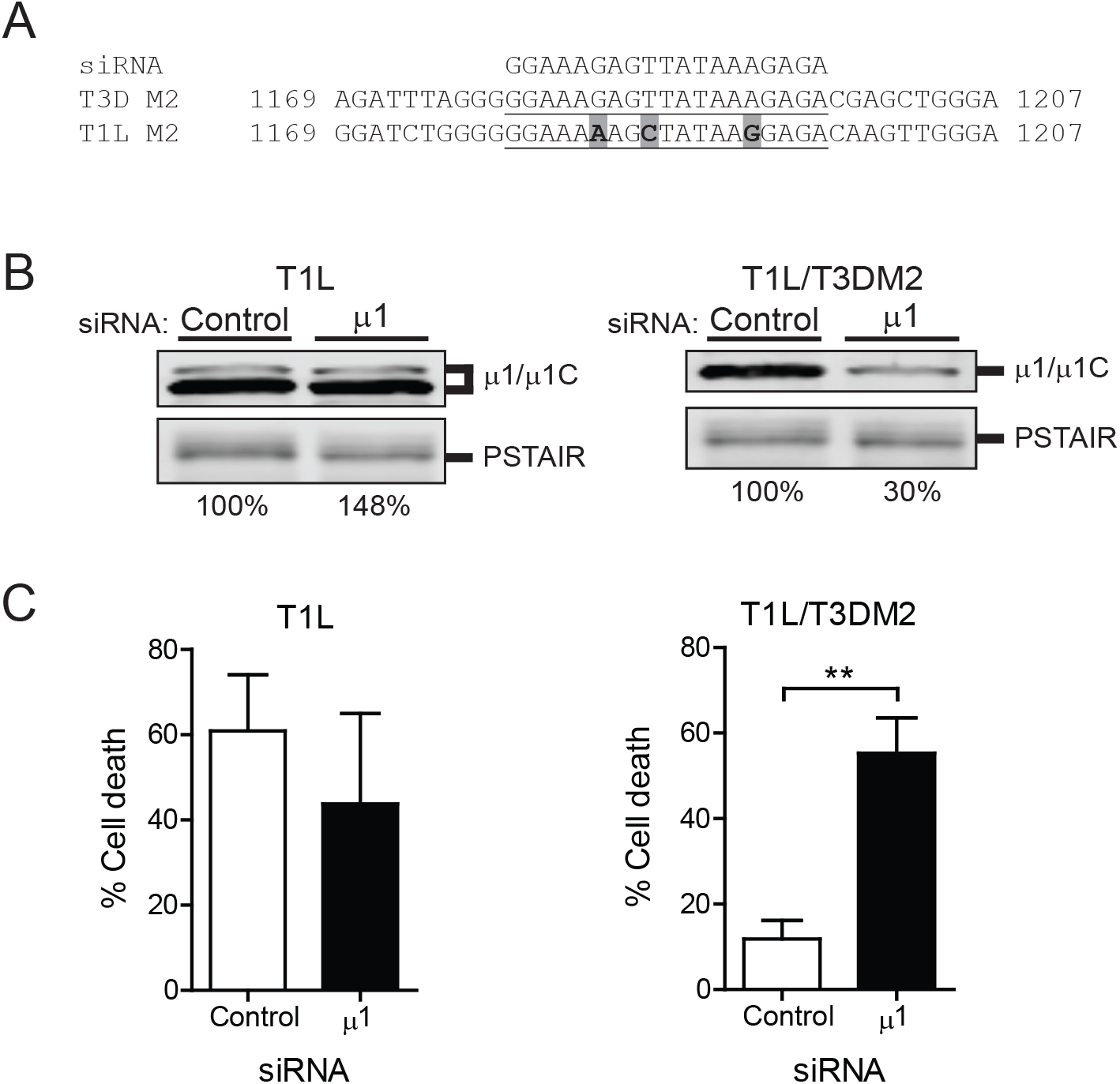
Effect of μ1 siRNA on cell death is sequence specific. (A) The μ1 siRNA sequence is aligned with the targeted region of the M2 gene segment from T3D. The corresponding region from T1L M2 showing mismatches within the siRNA target sequence is also shown. M2 encodes μ1. (B) L929 cells were transfected with either control β-gal or μ1 siRNA using INTERFERin. 24 h following transfection, the cells were infected with either T1L or T1L/T3DM2 at an MOI of 10 PFU/cell. Cell lysates prepared 24 h following infection were immunoblotted for the μ1 protein using anti-reovirus antisera and anti-PSTAIR mAb. The level of μ1 and μ1C relative to PSTAIR in control siRNA treated cells was considered 100%. (C) L929 cells were transfected with either control β-gal or μ1 siRNA using lipofectamine. 24 h following transfection, the cells were infected with either T1L or T1L/T3DM2 at an MOI of 10 PFU/cell for 72 h and 24 h respectively. Following infection, cell death was quantified by AOEB staining. Mean values for three independent infections are shown. Error bars indicate SD. P values were determined by student’s t-test. **, p < 0.005

To determine if μ1 knockdown is sufficient to influence virus replication, we measured viral yield over a 24 h interval. Because μ1 is an essential outer capsid protein whose function is required to launch infection (26), it is expected that an efficient knockdown of newly synthesized μ1 would result in a decrease in production of infectious progeny. We observed that knockdown of μ1 following T3D infection results in a significant (~ 1 log_10_) decrease in viral yield (Figure 4A). Since our siRNA targets newly synthesized μ1, it would not be expected that this knockdown would impact any events early in viral replication prior to gene expression. Consistent with this, we found that virus attachment and disassembly of incoming particles (which are derived from normal cells) are not affected in cells that express μ1 siRNA (not shown). Therefore, as expected, only events in viral replication that are dependent on *de novo* expression of μ1 impact viral yield.

**Figure 4.**
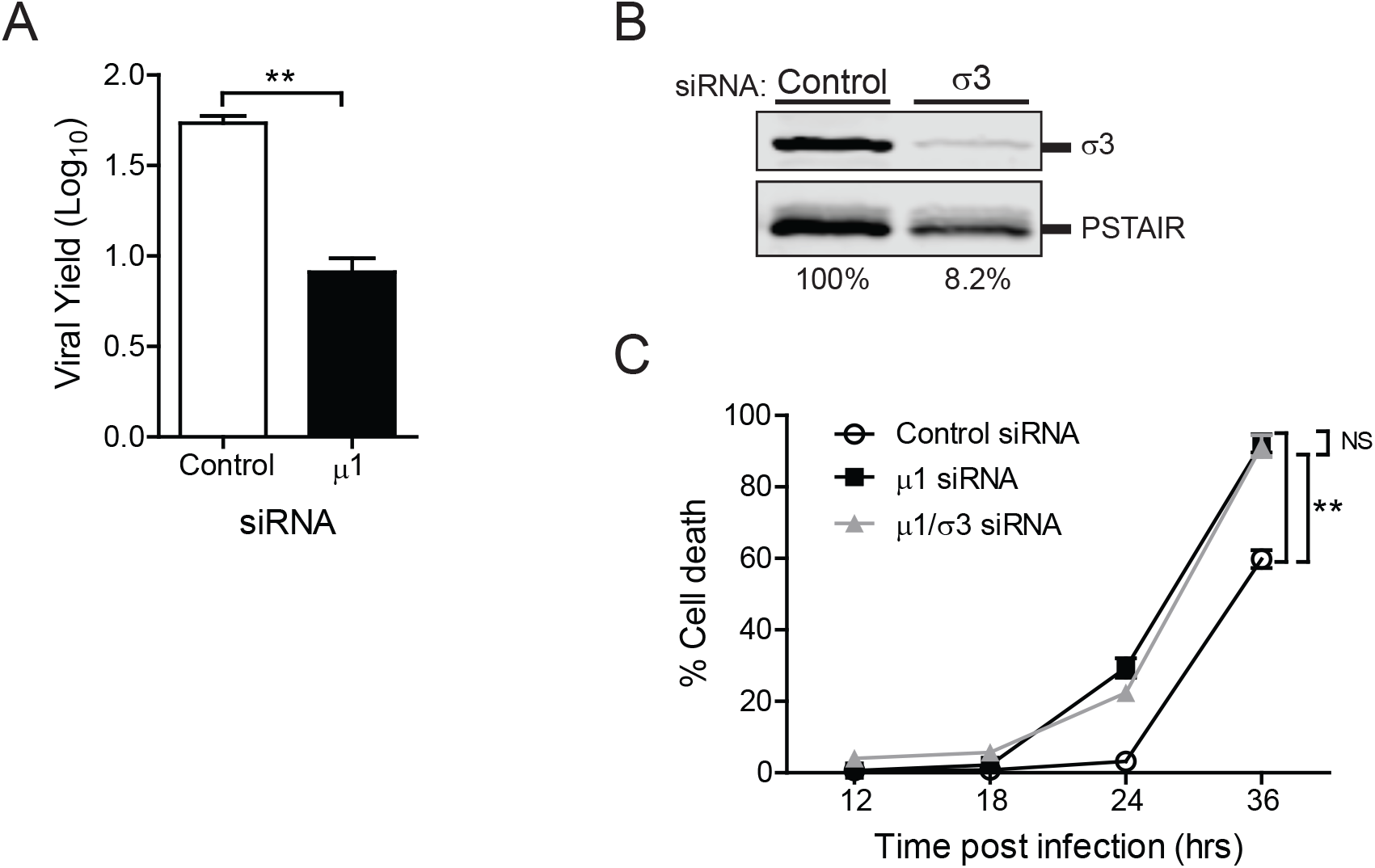
Knockdown of μ1 reduces viral yield and evokes cell death independently of σ3. (A) L929 cells were transfected with either control β-gal or μ1 siRNA using lipofectamine. 24 h following transfection, the cells were infected with T3D at an MOI of 2 PFU/cell. Viral replication 24 h following infection was measured by plaque assay on L929 cells. Mean viral yield for 3 independent infections are shown. (B) L929 cells were transfected with either control β-gal or σ3 siRNA using lipofectamine. 24 h following transfection, the cells were infected with T3D at an MOI of 10 PFU/cell. Cell lysates prepared 24 h following infection were immunoblotted for the σ3 protein using anti-reovirus antisera and anti-PSTAIR mAb. The level of σ3 relative to PSTAIR in control siRNA treated cells was considered 100%. (C) L929 cells were transfected with either control β-gal, μ1, or both μ1 and σ3 siRNA using INTERFERin. 24 h following transfection, the cells were infected with T3D at an MOI of 10 PFU/cell for 24 h. Following infection, cell death was quantified by AOEB staining. Mean values for three independent infections are shown. Error bars indicate SD. P values were determined by student’s t-test. **, p < 0.005

Newly synthesized μ1 interacts with its viral binding partner, σ3 (27-29). This interaction negatively impacts the function of μ1 in apoptotic cell death (12). Free σ3 promotes viral protein synthesis. However, when σ3 is complexed to μ1, the protein synthesis promoting function of σ3 is diminished (30, 31). To determine if the increase in cell death observed following μ1 knockdown is due to the presence of free σ3 in cells, L929 cells transfected with control or μ1 siRNA, or co-transfected with μ1 and σ3 siRNA (Figure 4B) were infected with T3D and cell death was quantified over a time course by AOEB staining (Figure 4C). Cells transfected with control siRNA-treated cells started exhibiting cell death around 36 h post infection with T3D. In contrast, cells transfected with μ1 siRNA succumbed to reovirus infection as early as 24 h post infection exhibiting cell death at ~30% as compared to control siRNA treated cells at ~5%. Moreover, in comparison to control siRNA transfected cells, significantly more cell death was observed at each time point in cells transfected with the μ1 siRNA. Knockdown of both μ1 and σ3 together demonstrated a phenotype similar to that of knocking down μ1 alone. In comparison to control siRNA treated cells, a greater level of cell death was also observed following knockdown of both μ1 and σ3. Moreover, cell death following double knockdown of μ1 and σ3 occurred with kinetics that resemble cell death following knockdown of only μ1. Thus, the enhancement of cell death following μ1 knockdown occurs regardless of whether σ3 is present in cells.

L929 cells infected with reovirus under control conditions undergo RIP3 dependent necroptosis (15, 16). It is possible, however, that μ1 knockdown changes the nature of cell death. To determine what form of cell death is enhanced by μ1 knockdown following T3D infection, we quantified cell death in μ1 knockdown cells treated with inhibitors of apoptotic or necroptotic cell death. Though treatment of cells with the pan-caspase inhibitor QVD completely blocked effector caspase (caspase-3/7) activity in reovirus infected cells (Figure 5A), it did not influence cell death in presence of μ1 siRNA knockdown (Figure 5B). These data suggest that following knockdown of μ1, reovirus continues to induce a non-apoptotic form of cell death. To block necroptosis, we used an siRNA specific to RIP3 (Figure 5C). Double knockdown of μ1 and RIP3 resulted in a diminishment of cell death (Figure 5D). Thus, even following knockdown of μ1, L929 cells infected with reovirus undergo cell death via necroptosis.

**Figure 5.**
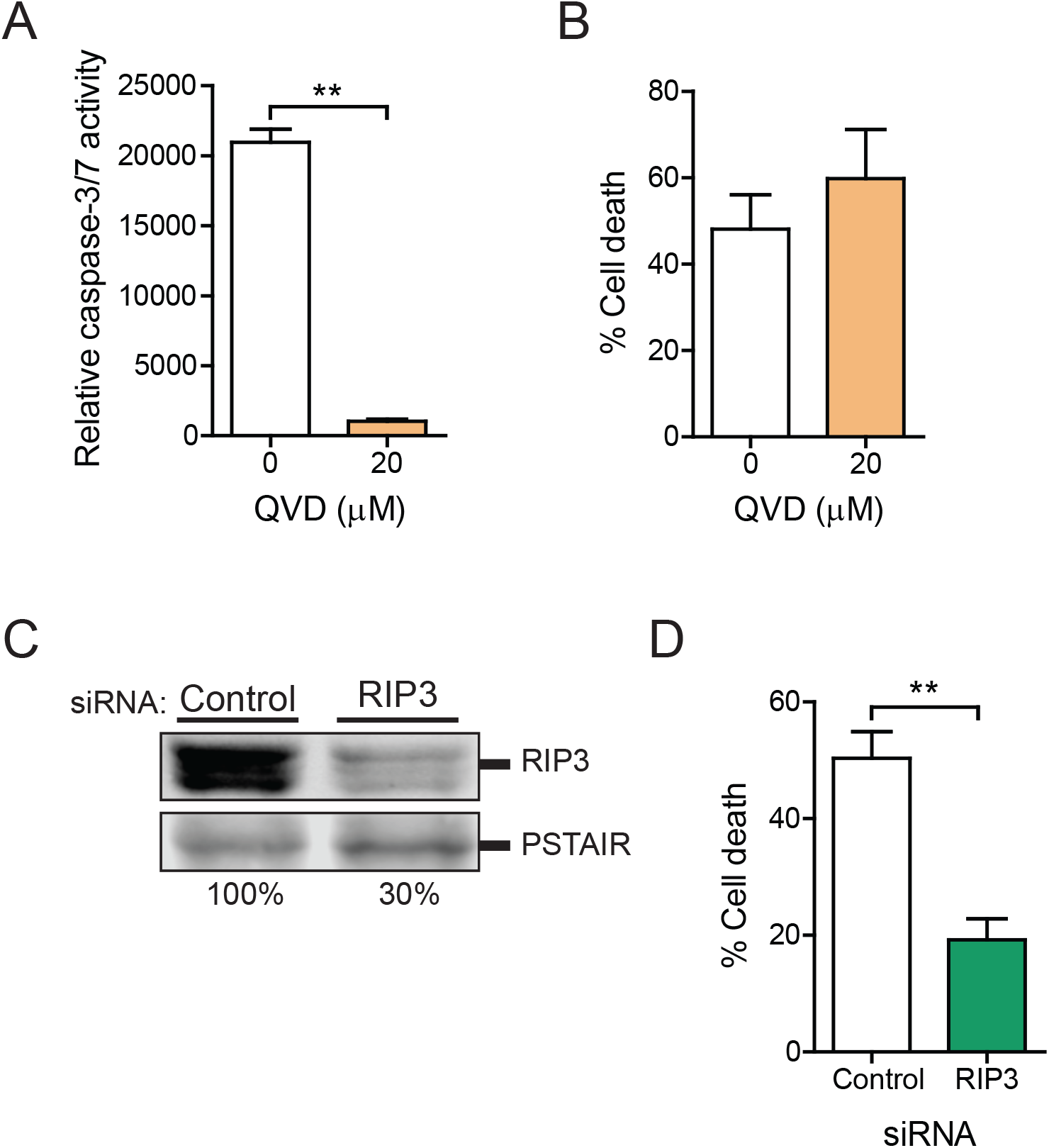
μ1 knockdown increases reovirus induced necroptosis in L929 cells. L929 cells were transfected with μ1 siRNA using lipofectamine. 24 h following transfection, the cells were infected with T3D at an MOI of 10 PFU/cell and either left untreated or were treated with 20 µM Q-VD-OPh when infection was initiated. At 30 h following infection, (A) caspase-3/7 activity was determined by chemiluminescent assay or (B) cell death was quantified by AOEB staining. (C) L929 cells were transfected with control β-gal or RIP3 siRNA using INTERFERin. Cell lysates prepared 24 h following transfection were immunoblotted using anti-RIP3 and anti-PSTAIR antibodies. The level of RIP3 relative to PSTAIR in control siRNA treated cells was considered 100%. (D) L929 cells were cotransfected with siRNA specific to μ1 and β-gal or μ1 and RIP3 using lipofectamine. 36 h following transfection cells were infected with T3D. At 30 h following infection cell death was quantified by AOEB staining. Mean values for three independent infections are shown. Error bars indicate SD. P values were determined by student’s t-test. **, p < 0.005

### Knockdown of μ1 increases reovirus-induced necroptosis downstream of IFN signaling

Based on previous data from our lab, we have proposed that necroptosis following reovirus infection requires expression of type I IFN and signaling via the IFNα/β receptor (IFNAR) to allow expression of a yet to be identified ISG required for cell death (16). Thus, one possible way in which μ1 knockdown increases cell death could be through an increase in the synthesis of type I IFN or through increased signaling via the IFNAR. We quantified levels of IFNβ mRNA and ZBP1 as a representative ISG in T3D-infected cells by RT-qPCR. At 18 h following infection, a similar increase in expression of IFNβ and ZBP1 mRNA was observed in control and μ1 siRNA treated cells (Figure 6A and 6B). These data suggest that enhanced induction of necroptosis in reovirus infected cells with a diminished level of μ1 is not related to enhanced IFNβ induction or IFNAR signaling.

**Figure 6.**
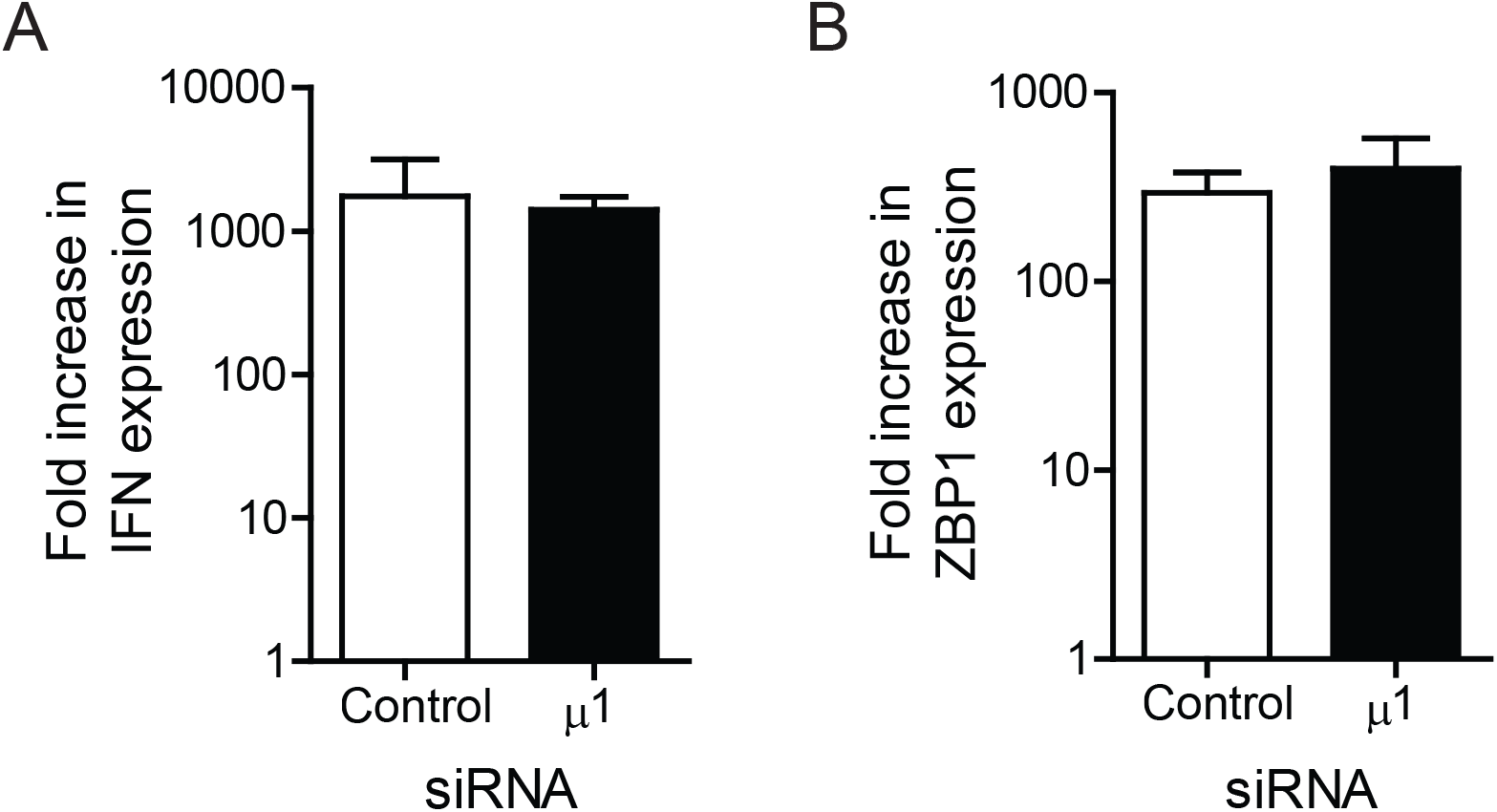
Reduction in μ1 levels does not impact IFN signaling following infection. L929 cells were transfected with β-gal or μ1 siRNA using INTERFERin. 24 h following transfection, the cells were infected with T3D at an MOI of 10 PFU/cell. RNA extracted from infected cells harvested at 0 and 18 h post infection was reverse transcribed using random primers. Fold increase in levels of (A) IFNβ or (B) ZBP1 mRNA relative to GAPDH mRNA over 18 h of infection was quantified by qPCR and comparative CT analyses. Mean values for three independent infections are shown. Error bars indicate SD.

### Viral RNA genome and secondary transcripts accumulate following μ1 knockdown

Our previous evidence indicated that in addition to IFN signaling, *de novo* generation of viral dsRNA genome and/or a product dependent on dsRNA synthesis contributes to the induction of necroptosis by reovirus (16). Thus, one way in which μ1 knockdown can result in greater cell death is due to an increased accumulation of these viral components in infected cells. To address this question, we measured the levels of viral minus and plus strand RNA in infected cells by strand-specific reverse transcription and qPCR. We used S1 gene derived RNAs as representatives for these experiments. In comparison to control siRNA treated cells, μ1 siRNA treated cells contained ~4 fold more minus strand RNA at 21 h post infection (Figure 7A). Since minus strand RNA only exists in the context of dsRNA genome during reovirus replication (32), minus strand levels are indicative of the amount of genomic dsRNA. Thus, μ1 knockdown increases the level of progeny dsRNA. A significantly greater (~2-3 fold) level of S1 plus strand in cells was also observed in cells transfected with μ1 siRNA (Figure 7B). While we observed an increase in accumulation of both viral minus and plus strand RNA following knockdown of μ1, under both control and μ1 knockdown conditions there is a substantially larger quantity (~100 fold) more plus strand RNA relative to minus strand. Furthermore, there is no significant difference in the levels of plus strand RNA relative to minus strand RNA when μ1 is knocked down as compared to control siRNA-treated cells (Figure 7C).

**Figure 7.**
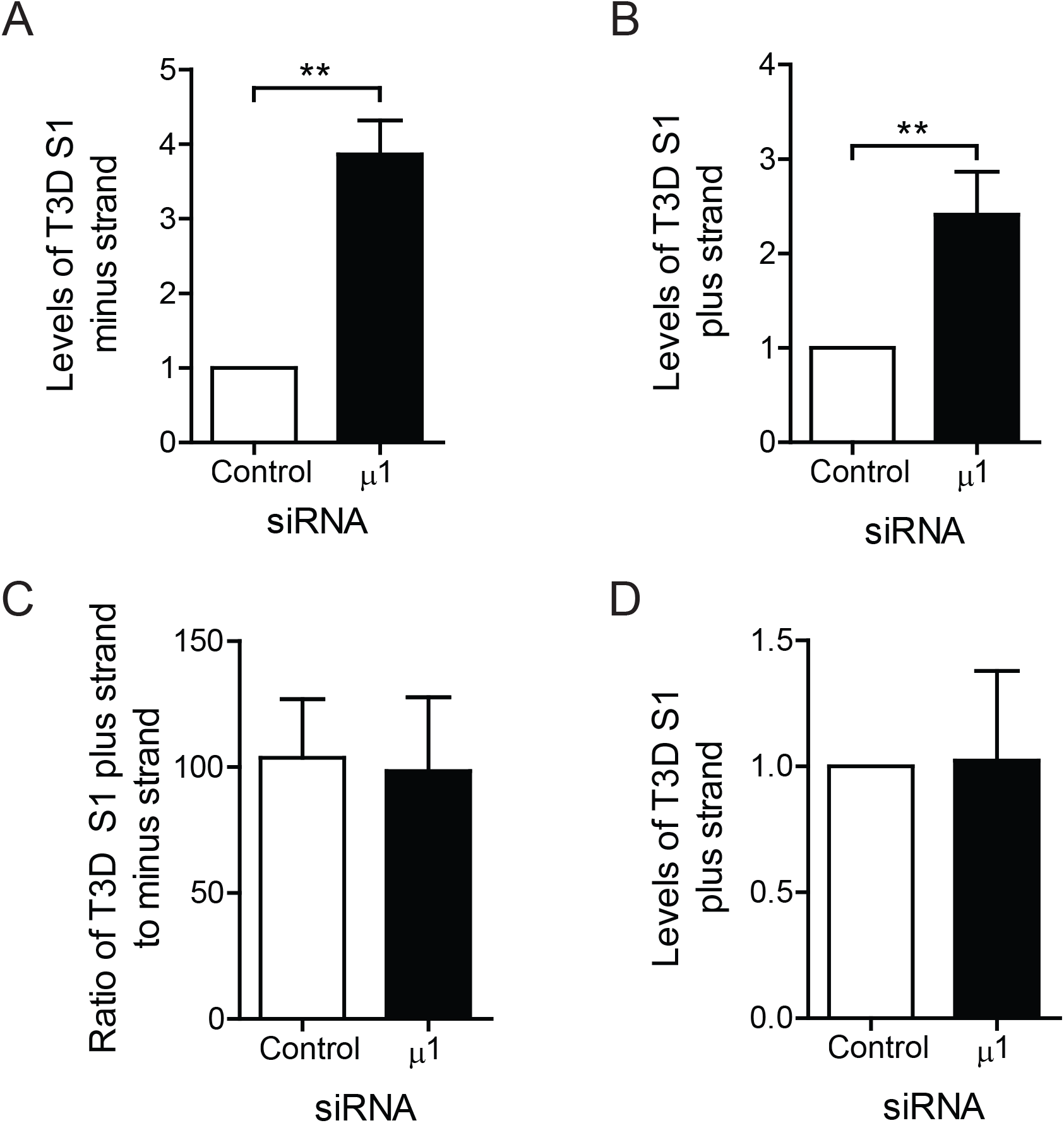
μ1 knockdown results in increased accumulation of viral RNAs following infection. L929 cells were transfected with control β-gal or μ1 siRNA using INTERFERin. 24 h following transfection, cells were infected with T3D at an MOI of 10 PFU/cell. RNA extracted from (A,B,C) untreated infected cells harvested at 21 h post infection, or (D) cells treated with 30mM GuHCl at time of infection harvested at 8 h post infection, was reverse transcribed using primers specific to the minus strand or plus strand of the T3D S1 gene or GAPDH mRNA. Levels of accumulated T3D S1 (A) minus strand or (B and D) plus strand RNA relative to GAPDH mRNA was quantified by qPCR and comparative C_T_ analysis. (A, B, and D) The ratio of T3D S1 RNA to GAPDH in control siRNA-treated cells was set to 1. (C) The level of plus strand relative to minus strand was quantified by qPCR and comparative C_T_ analysis. Mean values for three independent infections are shown. Error bars indicate SD. P values were determined by student’s t-test. **, p < 0.005

In reovirus infected cells, plus strand RNA is present in the context of viral genomic dsRNA and in the form of viral primary and secondary transcripts (33-36). Since the experiment above was performed at 21 h, our plus strand measurement represents a combination of each of these plus strand species. Under the conditions used, plus strand RNA of the incoming genome is near the lower limit of detection of this assay and does not impact our measurement (data not shown). It is expected that if plus strand RNA is quantified early in infection, it would predominantly represent primary transcripts. Inclusion of GuHCl, which blocks dsRNA synthesis and therefore secondary transcription (37), would further ensure that only primary transcripts are present in cells. To determine whether μ1 knockdown affects primary transcription, we measured viral plus strand level at 8 h post infection in the presence of GuHCl. Under these conditions, the S1 plus strand accumulated to equivalent levels in control and μ1 siRNA treated cells (Figure 7D). Because primary transcript levels are unchanged following μ1 knockdown, these data suggest that the increase in plus strand RNA observed later in infection is related either to an increase in viral dsRNA, secondary transcripts, or both.

An increase in secondary mRNA transcripts should also result in an increase in the accumulation of viral proteins (38-42). In comparison to control cells, in cells transfected with μ1 siRNA, we observed a significant increase in two representative viral proteins - σNS and σ3 (Figure 8A and 8B). Together our data suggest that products of reovirus infection – genomic dsRNA, secondary transcripts, and viral proteins – accumulate to higher levels when μ1 is knocked down.

**Figure 8.**
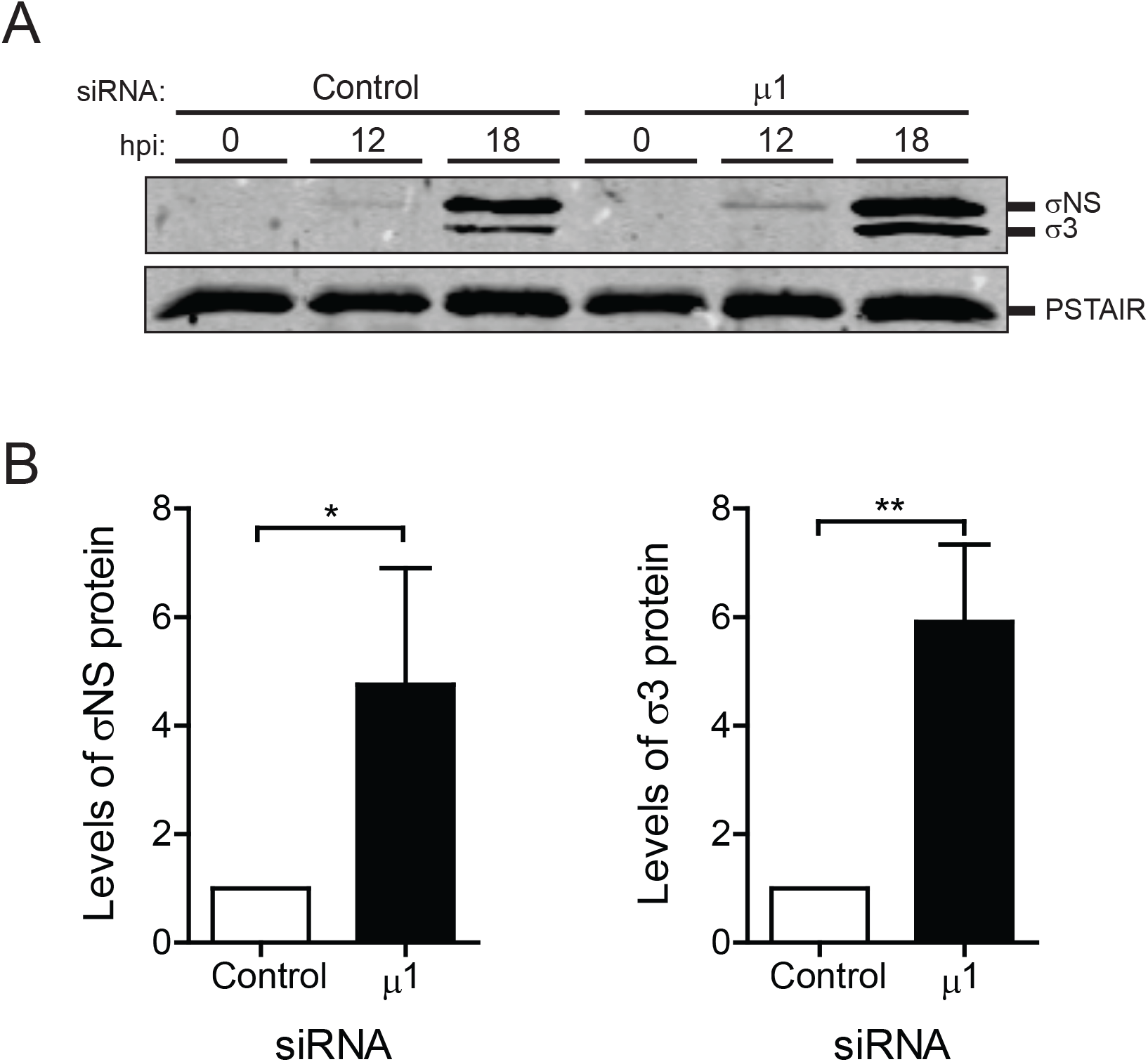
Knockdown of μ1 increases accumulation of other viral proteins following infection. L929 cells were transfected with control β-gal or μ1 siRNA using lipofectamine. 24 h following transfection, cells were infected with T3D at an MOI of 10 PFU/cell. (A) Cell lysates prepared 0, 12, or 18 h following infection were immunoblotted for the σNS protein with anti-σNS antisera, σ3 protein with anti-reovirus antisera, and anti-PSTAIR mAb. (B) Quantification of accumulated levels of σNS (left) and σ3 (right) relative to PSTAIR at 18 h post infection are shown as mean values from three independent infections. The ratio of viral protein to GAPDH for control siRNA-treated cells was set to 1. Error bars indicate SD. P values were determined by student’s t-test. *,p < 0.05; **,p < 0.005

### Knockdown of μ1 results in a decreased sensitivity to the blocking effects of GuHCl

Treatment of reovirus infected cells with GuHCl results in blockade of progeny dsRNA synthesis and a concomitant reduction in necroptotic cell death (16, 37). Because we observed a greater level of dsRNA in infected cells following μ1 knockdown (Fig. 7), we asked if the potency with which GuHCl influences dsRNA synthesis is altered in the absence of μ1. We found that at 21 h post infection in control siRNA treated cells, GuHCl treatment diminishes levels of the S1 minus strand RNA in a dose-dependent manner. Treatment with 15 mM GuHCl was sufficient to reduce levels of S1 minus strand by about 60% (Fig 8A). In contrast, 15 mM GuHCl did not affect levels of S1 minus strand RNA levels in cells transfected with μ1 siRNA. Thus, μ1 knockdown influences the sensitivity of genomic dsRNA synthesis to GuHCl. We also measured the impact of GuHCl on the levels of viral plus strand RNA following μ1 knockdown by RT-qPCR (Figure 9B). Similar to what was observed for minus strand RNA levels, at 21 h post infection in control siRNA treated cells, GuHCl treatment reduced levels of S1 plus strand RNA in a dose-dependent manner under control conditions. Treatment of control siRNA-treated cells with 15 mM GuHCl reduced levels of S1 plus strand RNA by ~60%. In contrast, 15 mM GuHCl reduced levels of S1 plus strand RNA by only ~25% following μ1 knockdown. Therefore, μ1 knockdown also decreases the sensitivity of viral plus strand RNA to GuHCl. Consistent with μ1 knockdown causing an increased accumulation of these viral gene products, there is significantly more minus and plus strand RNA accumulated in μ1 siRNA-treated cells relative to control siRNA-treated cells in the presence of each concentration of GuHCl (not shown).

**Figure 9.**
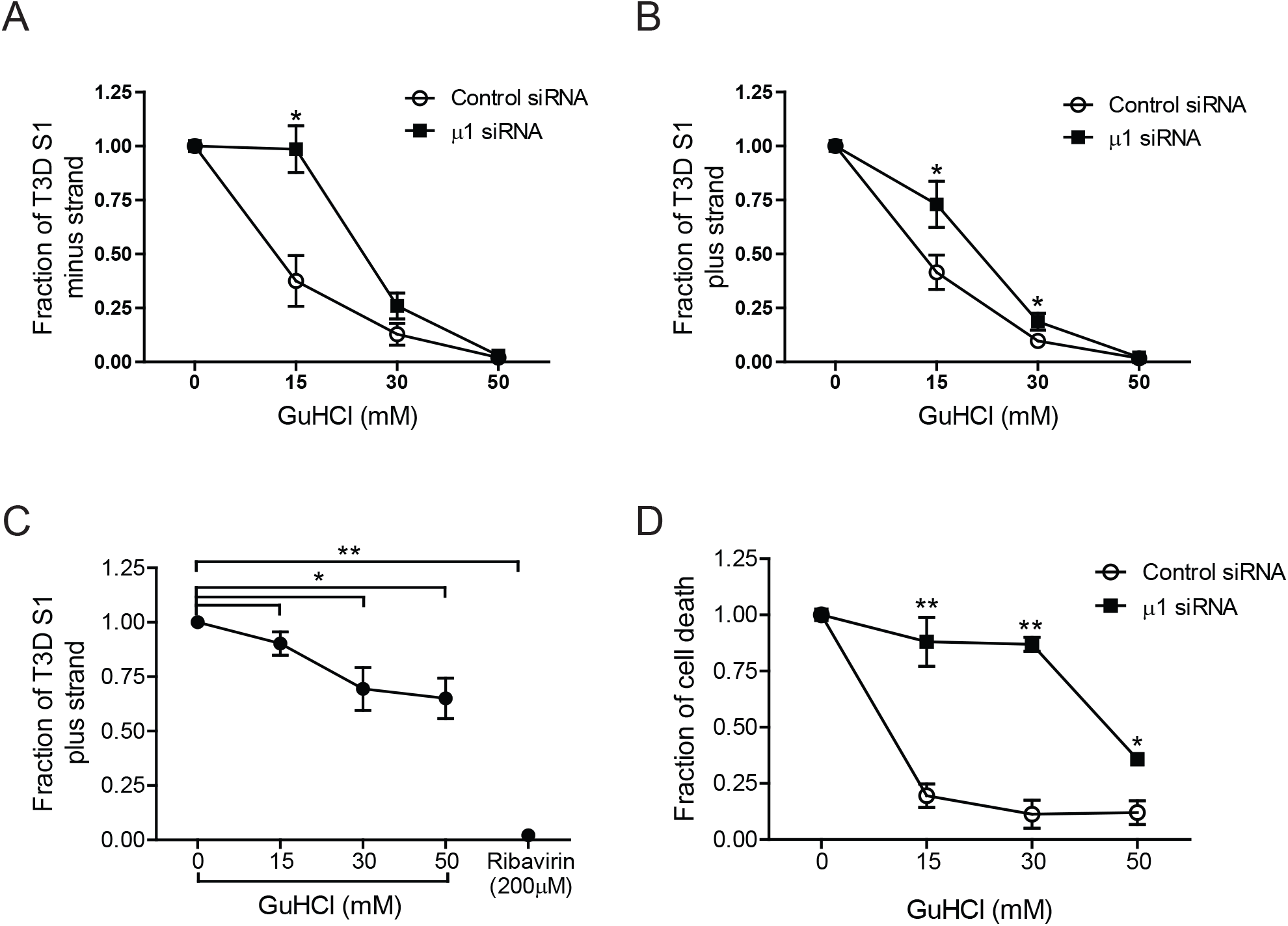
Intracellular μ1 levels affect sensitivity of viral RNA synthesis and cell death to GuHCl. L929 cells were transfected with control β-gal or μ1 siRNA using INTERFERin. 24 h following transfection cells were infected with T3D at an MOI of 10 PFU/cell and processed using conditions described. (A-D) Cells were either left untreated, or 200 µM ribavirin or GuHCl at the indicated concentrations was added when infection was initiated. RNA extracted from cells harvested (A and B) 21 h or (C) 6 h following infection was reverse transcribed using primers specific to the minus strand or plus strand of the T3D S1 gene or GAPDH mRNA. Fraction of accumulated T3D S1 (A) minus strand or (B and C) plus strand relative to GAPDH mRNA in the presence of GuHCl or ribavirin was quantified by qPCR and comparative C_T_ analysis for three independent infections. T3D S1 accumulation in the absence of treatment was set to 1 for each siRNA-treated sample. (D) At 30 h post infection, cell death was quantified by AOEB staining. Fraction of cell death in the presence of GuHCl was quantified for three independent infections. Cell death in the absence of GuHCl was set to 1 for each siRNA-treated sample. Error bars indicated SD. P values were determined by Student’s t-test. *, p < 0.05; **, p < 0.005

Previously, studies have only used concentrations up to 15 mM GuHCl to investigate plus and minus synthesis following reovirus infection (16, 37). Because we used a significantly higher concentration of GuHCl, we sought to confirm that GuHCl does not impact primary transcription. Toward this end, we quantified levels of S1 plus strand by RT-qPCR early at 6 h post infection in infected cells (Figure 9C). Ribavirin, which potently inhibits primary rounds of plus strand synthesis (43), was used as a control. As expected, ribavirin caused a near complete inhibition of the synthesis of plus strand RNA. Though treatment with increasing concentration of GuHCl resulted in a reduction in S1 plus strand levels, this reduction was significantly modest. Even at 50 mM GuHCl, no greater than ~ 30% reduction in the levels of plus strand RNA was observed. We did not directly assess how high concentration of GuHCl slightly inhibits plus strand RNA synthesis early in infection. However, based on a significantly greater of reduction of plus strand RNA later in infection, we conclude that GuHCl mainly impacts later stages of infection that are dependent on minus strand RNA synthesis, such as secondary transcription.

To determine if knockdown of μ1 also confers a decrease in sensitivity of cell death to GuHCl, we transfected cells with either control or μ1 siRNA and infected with T3D in the presence of increasing concentrations of GuHCl. At 30 h post infection, cell death was quantified by AOEB staining (Figure 9D). Similar to the effect of GuHCl on RNA levels we observed, control siRNA treated cells demonstrated a dose dependent decrease in cell death. 15 mM GuHCl was sufficient to reduce cell death by ~75%. In contrast, when μ1 was knocked down, even a significantly higher concentration of GuHCl (50 mM) was only able to reduce cell death by ~ 50%. Together, the decreased sensitivity of minus strand synthesis, secondary transcription and cell death to GuHCl following μ1 knockdown suggests that the increase in cell death observed may be due to an increase in one or more viral products.

## DISCUSSION

In this study, we assessed the role of newly synthesized μ1 in cell death following reovirus infection. We report that whereas knockdown of μ1 does not impact apoptosis induced by reovirus, μ1 knockdown enhances necroptotic cell death following reovirus infection. The loss of μ1 is accompanied by an increase in accumulation of viral minus strand RNA (a measure of dsRNA), plus strand RNA (which are likely secondary transcripts produced from newly synthesized dsRNA), and viral proteins. Moreover, cell death following μ1 knockdown was less sensitive to GuHCl, an agent that blocks dsRNA synthesis. Based on these data we conclude that increased accumulation of one of these viral gene products in the absence of μ1 results in enhanced cell death following reovirus infection. These data raise the following three questions:(i) how does μ1 knockdown affect the generation of viral gene products; (ii) how do μ1 levels influence sensitivity of virus replication to GuHCl; and (iii) how does an increase in viral gene products correlate with enhanced cell death by necroptosis. These issues are discussed below.

An intriguing observation from our experiments is the increased accumulation of viral products in absence of μ1. Viral RNA is transcribed at two stages of reovirus replication: (i) during entry, by cores formed from incoming viral particles to generate primary transcripts and, during assembly, by progeny cores to generate secondary transcripts (33-36). Transcriptional activity following entry of the virus into cells correlates with loss of the outer capsid. In vitro studies demonstrate that cores, which lack μ1, and ISVP*s, which contain μ1 in an altered conformation, are transcriptionally active (44-48). In contrast, virions and ISVPs, which contain μ1 in its native conformation are not. When μ1 is in its native conformation, such as in virions and ISVPs, λ2, a core protein that forms turrets at the 12 icosahedral vertices of the virus, assumes a closed conformation (49). In contrast, λ2 assumes an open conformation in cores (49, 50). An open conformation for λ2 correlates with transcriptional activity, potentially because this allows for entry of nucleotides and the exit of viral mRNA from these turrets (49, 50). Structural information on λ2 in ISVP*s is lacking but because ISVP*s are capable of transcription, it is likely that λ2 conformation in ISVP*s more resembles that in cores than in ISVPs. How transcriptional activity of progeny cores is regulated is not understood. However, based on the regulation of transcription during entry, it is reasonable to hypothesize that assembly of the outer capsid comprised of μ1-σ3 heterohexamers onto progeny cores closes the structure of λ2, thereby shutting off transcription. We reason that when μ1 is knocked down, outer capsid assembly is rendered inefficient. Thus, an increased proportion of progeny cores are capable of performing secondary rounds of transcription or that secondary transcription proceeds for a more extended period (Figure 10). We think that this may explain the increase in viral plus strand RNA accumulation later in infection. Because the plus strand RNA can be translated, we also observe an increase in the level of viral proteins. Finally, because the plus strand is packaged and used as a template for minus strand synthesis, we also observe a greater level of minus strand RNA in absence of μ1. We hypothesize that continued propagation of this replication loop results in the accumulation of viral mRNA, protein, and minus strand RNA we observed following loss of μ1.

**Figure 10.**
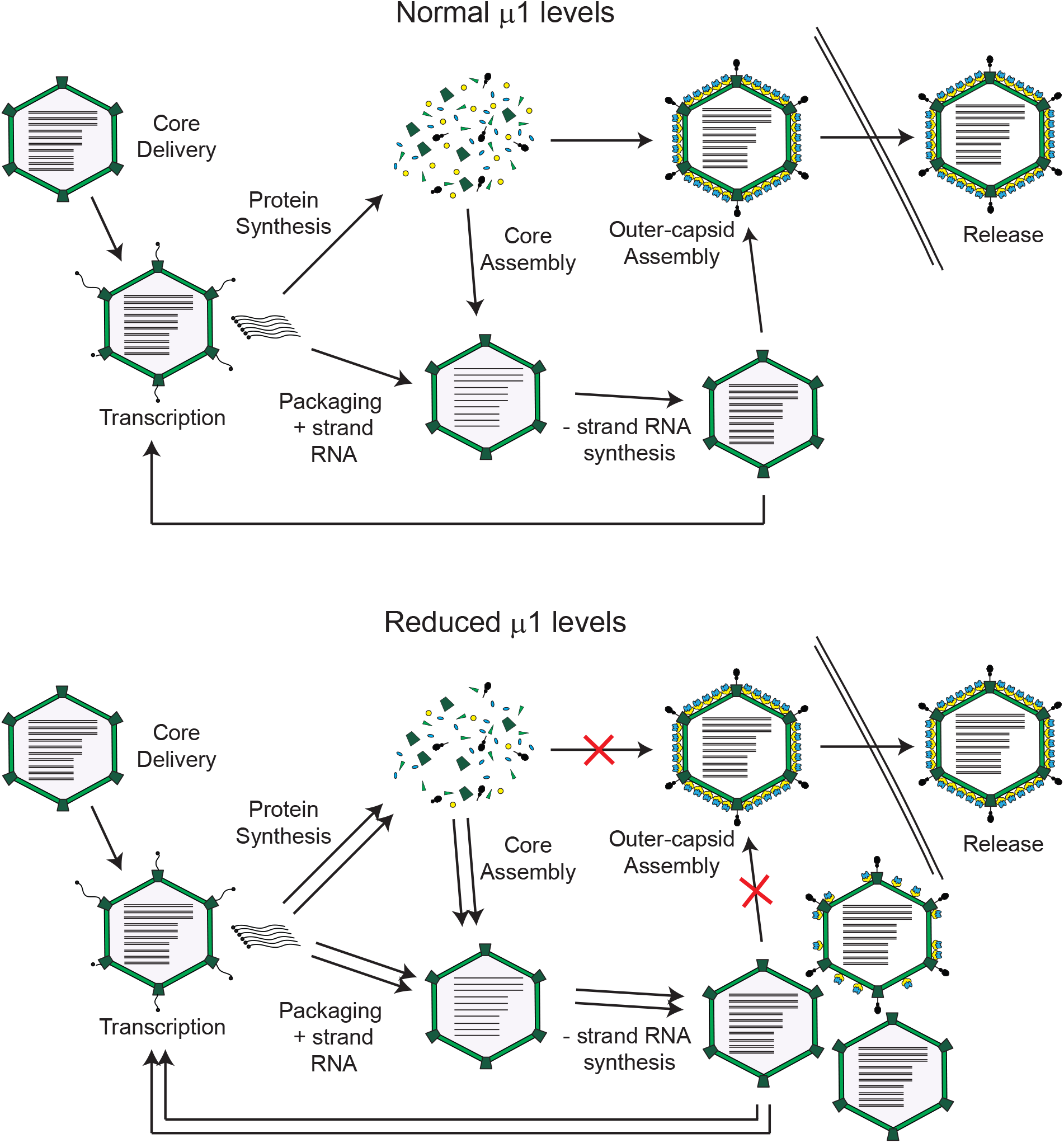
Model depicting late stages of reovirus replication following knockdown of μ1. Stages of reovirus replication proposed to be enhanced following knockdown of μ1 are indicated by double arrows.

In reovirus infected cells, GuHCl blocks progeny viral dsRNA synthesis (37). Because secondary transcription is dependent on dsRNA synthesis, GuHCl also diminishes secondary transcription. However, the viral target of GuHCl is not known and the basis for how GuHCl blocks these events during reovirus replication is not understood. Since GuHCl affects the synthesis of new dsRNA, it is possible that the viral target is the polymerase itself. However, because GuHCl has no effect on primary rounds of transcription, this would suggest that the polymerase functions differently during plus and minus strand RNA synthesis. Alternatively, GuHCl may target a factor which associates with progeny cores during dsRNA synthesis. Such a factor could include a viral nonstructural protein or an outer capsid protein that is known to interact with progeny cores, such as μ1. We observed that in the absence of μ1, GuHCl is less capable of inhibiting viral dsRNA synthesis and secondary transcription. Thus, our data support the possibility that μ1 is the target of GuHCl mediated effects on viral replication. However, direct evidence for this idea is thus far lacking.

Our previous work has demonstrated a role for newly synthesized viral dsRNA in the induction of necroptosis by reovirus. Our data presented here suggest that necroptosis induction may require viral dsRNA, or products dependent on its synthesis such as secondary transcripts or maximal level of viral proteins (figures 7-9). Genomic dsRNA is generated within core particles which are embedded in a viral factory matrix produced by viral non-structural proteins. This arrangement is thought to prevent host detection of viral genomic RNA in infected cells. Nonetheless, recent evidence indicates that reovirus factories can be stained by a dsRNA specific antibody (51), suggesting that at least proportion of dsRNA is accessible and likely available for detection. If this accessibility is altered by μ1 knockdown, it could result in enhanced cell death. During primary rounds of transcription of reovirus RNA, the synthesized messages are capped and utilized for translation or packaging (52, 53). However, secondary transcripts, synthesized late in infection, are uncapped (36). If uncapped transcripts are the viral trigger for necroptosis, it is possible that an increase in secondary transcripts produced by μ1 knockdown therefore increases the amount of viral signature in infected cells, thereby leading to enhanced necroptosis. Nonetheless, potential cellular sensors of dsRNA or secondary transcripts that contribute to cell death are yet to be identified. The loss of μ1 also results in an increase in levels of other viral proteins (Figure 8). Thus, increased levels of one or more viral proteins may also result in cell death. While increase in σ3 produced by μ1 knockdown does not contribute to necroptosis, the accumulation of other viral proteins may also lead to enhanced necroptosis. Connecting viral components to initiate signaling that leads to necroptosis requires further investigation.

In conclusion, this work identifies a new role for the viral protein μ1 in limiting the transcriptional activity of newly synthesized cores. Furthermore, the loss of μ1-mediated regulation of viral replication events late in infection results in enhanced cell death induction. These intriguing data also highlight that in addition to μ1 playing an essential role in viral entry, μ1 can impact two different cell death pathways. μ1 from the incoming viral capsid potentiates the activation of proapoptotic signaling pathways (10, 11). In contrast, newly synthesized μ1 that is generated in infected cells indirectly, by decreasing the accumulation of viral gene products, diminishes the levels of necroptosis following reovirus. Reovirus is currently being tested as an oncolytic agent due to its ability to preferentially replicate in, and kill, cancer cells. Thus, optimizing the capacity for reovirus to induce a specific form of cell death by manipulation of the properties of the μ1 protein could be highly advantageous for maximizing its oncolytic potential.

## ACKNOWLEDGEMENTS

We thank members of our laboratory and the Indiana University virology community for helpful suggestions.

Research reported in this publication was supported by the National Institute of Allergy and Infectious Diseases of the National Institutes of Health under award number 1R01AI110637 (to P.D.). The content is solely the responsibility of the authors and does not necessarily represent the official views of the funders.

## REFERENCES

1. Danthi P. 2016. Viruses and the Diversity of Cell Death. Annu Rev Virol 3:533–553.

2. Upton JW, Chan FK. 2014. Staying Alive: Cell Death in Antiviral Immunity. Mol Cell 54:273–280.

3. DeBiasi RL, Robinson BA, Sherry B, Bouchard R, Brown RD, Rizeq M, Long C, Tyler KL. 2004. Caspase inhibition protects against reovirus-induced myocardial injury in vitro and in vivo. Journal of Virology 78:11040–11050.

4. Oberhaus SM, Smith RL, Clayton GH, Dermody TS, Tyler KL. 1997. Reovirus infection and tissue injury in the mouse central nervous system are associated with apoptosis. J Virol 71:2100–6.

5. Danthi P, Pruijssers AJ, Berger AK, Holm GH, Zinkel SS, Dermody TS. 2010. Bid regulates the pathogenesis of neurotropic reovirus. PLoS Pathog 6:e1000980.

6. Tyler KL, Squier MK, Rodgers SE, Schneider BE, Oberhaus SM, Grdina TA, Cohen JJ, Dermody TS. 1995. Differences in the capacity of reovirus strains to induce apoptosis are determined by the viral attachment protein sigma 1. J Virol 69:6972–9.

7. Tyler KL, Squier MK, Brown AL, Pike B, Willis D, Oberhaus SM, Dermody TS, Cohen JJ. 1996. Linkage between reovirus-induced apoptosis and inhibition of cellular DNA synthesis: role of the S1 and M2 genes. J Virol 70:7984–91.

8. Connolly JL, Barton ES, Dermody TS. 2001. Reovirus binding to cell surface sialic acid potentiates virus-induced apoptosis. J Virol 75:4029–39.

9. Danthi P, Hansberger MW, Campbell JA, Forrest JC, Dermody TS. 2006. JAM-A-independent, antibody-mediated uptake of reovirus into cells leads to apoptosis. J Virol 80:1261–70.

10. Danthi P, Coffey CM, Parker JS, Abel TW, Dermody TS. 2008. Independent regulation of reovirus membrane penetration and apoptosis by the mu1 phi domain. PLoS Pathog 4:e1000248.

11. Danthi P, Kobayashi T, Holm GH, Hansberger MW, Abel TW, Dermody TS. 2008. Reovirus apoptosis and virulence are regulated by host cell membrane penetration efficiency. J Virol 82:161–72.

12. Coffey CM, Sheh A, Kim IS, Chandran K, Nibert ML, Parker JS. 2006. Reovirus outer capsid protein m1 induces apoptosis and associates with lipid droplets, endoplasmic reticulum, and mitochondria. Journal of Virology 80:8422–8438.

13. Wisniewski ML, Werner BG, Hom LG, Anguish LJ, Coffey CM, Parker JS. 2011. Reovirus infection or ectopic expression of outer capsid protein micro1 induces apoptosis independently of the cellular proapoptotic proteins Bax and Bak. J Virol 85:296–304.

14. Connolly JL, Dermody TS. 2002. Virion disassembly is required for apoptosis induced by reovirus. J Virol 76:1632–41.

15. Berger AK, Danthi P. 2013. Reovirus activates a caspase-independent cell death pathway. mBio 4.

16. Berger AK, Hiller BE, Thete D, Snyder AJ, Perez E, Jr., Upton JW, Danthi P. 2017. Viral RNA at Two Stages of Reovirus Infection Is Required for the Induction of Necroptosis. J Virol 91.

17. Newton K, Manning G. 2016. Necroptosis and Inflammation. Annu Rev Biochem 85:743–63.

18. Hiller BE, Berger AK, Danthi P. 2015. Viral gene expression potentiates reovirus-induced necrosis. Virology doi:10.1016/j.virol.2015.06.018.

19. Becker MM, Peters TR, Dermody TS. 2003. Reovirus sigma NS and mu NS proteins form cytoplasmic inclusion structures in the absence of viral infection. J Virol 77:5948–63.

20. Kobayashi T, Ooms LS, Ikizler M, Chappell JD, Dermody TS. 2010. An improved reverse genetics system for mammalian orthoreoviruses. Virology 398:194–200.

21. Berard A, Coombs KM. 2009. Mammalian reoviruses: propagation, quantification, and storage. Curr Protoc Microbiol Chapter 15:Unit15C 1.

22. Middleton JK, Severson TF, Chandran K, Gillian AL, Yin J, Nibert ML. 2002. Thermostability of reovirus disassembly intermediates (ISVPs) correlates with genetic, biochemical, and thermodynamic properties of major surface protein mu1. J Virol 76:1051–61.

23. Schmittgen TD, Livak KJ. 2008. Analyzing real-time PCR data by the comparative C(T) method. Nat Protoc 3:1101–8.

24. Kim JW, Lyi SM, Parrish CR, Parker JS. 2011. A proapoptotic peptide derived from reovirus outer capsid protein {micro}1 has membrane-destabilizing activity. Journal of virology 85:1507–16.

25. Connolly JL, Rodgers SE, Clarke P, Ballard DW, Kerr LD, Tyler KL, Dermody TS. 2000. Reovirus-induced apoptosis requires activation of transcription factor NF-kappaB. J Virol 74:2981–9.

26. Danthi P, Holm G. H., Stehle T., and Dermody T.S.. 2013. Reovirus receptors, cell entry, and signaling. *In* Pöhlmann S, and Simmons G. (ed), Viral Entry into Cells, Georgetown, TX.

27. Tillotson L, Shatkin AJ. 1992. Reovirus polypeptide s3 and N-terminal myristoylation of polypeptide m1 are required for site-specific cleavage to m1C in transfected cells. Journal of Virology 66:2180–2186.

28. Shepard DA, Ehnstrom JG, Schiff LA. 1995. Association of reovirus outer capsid proteins s3 and m1 causes a conformational change that renders s3 protease sensitive. Journal of Virology 69:8180–8184.

29. Liemann S, Chandran K, Baker TS, Nibert ML, Harrison SC. 2002. Structure of the reovirus membrane-penetration protein, m1, in a complex with its protector protein, s3. Cell 108:283–295.

30. Schmechel S, Chute M, Skinner P, Anderson R, Schiff L. 1997. Preferential translation of reovirus mRNA by a s3-dependent mechanism. Virology 232:62–73.

31. Yue Z, Shatkin AJ. 1997. Double-stranded RNA-dependent protein kinase (PKR) is regulated by reovirus structural proteins. Virology 234:364–371.

32. Sakuma S, Watanabe Y. 1972. Reovirus replicase-directed synthesis of double-stranded ribonucleic acid. Journal of Virology 10:628–638.

33. Sakuma S, Watanabe Y. 1971. Unilateral synthesis of reovirus double-stranded ribonucleic acid by a cell free replicase system. Journal of Virology 8:190–196.

34. Skup D, Millward S. 1980. mRNA capping enzymes are masked in reovirus progeny subviral particles. Journal of Virology 34:490–496.

35. Morgan EM, Zweerink HJ. 1975. Characterization of transcriptase and replicase particles isolated from reovirus infected cells. Virology 68:455–466.

36. Zarbl H, Skup S, Millward S. 1980. Reovirus progeny subviral particles synthesize uncapped mRNA. Journal of Virology 34:497–505.

37. Murray KE, Nibert ML. 2007. Guanidine hydrochloride inhibits mammalian orthoreovirus growth by reversibly blocking the synthesis of double-stranded RNA. J Virol 81:4572–84.

38. Skup D, Millward S. 1980. Reovirus-induced modification of cap dependent translation in infected L cells. Proceedings of the National Academy of Sciences USA 77:152–156.

39. Skup D, Zarbl H, Millward S. 1981. Regulation of translation in L-cells infected with reovirus. Journal of Molecular Biology 151:35–55.

40. Lemieux R, Zarbl H, Millward S. 1984. mRNA discrimination in extracts from uninfected and reovirus-infected L-cells. Journal of Virology 51:215–222.

41. Sonenberg N, Skup D, Trachsel H, Millward S. 1981. In vitro translation in reovirus- and poliovirus-infected cell extracts. Effects of anti-cap binding protein monoclonal antibody. J Biol Chem 256:4138–41.

42. Detjen BM, Walden WE, Thach RE. 1982. Translational specificity in reovirus-infected mouse fibroblasts. Journal of Biological Chemistry 257:9855–9860.

43. Rankin UT, Jr., Eppes SB, Antczak JB, Joklik WK. 1989. Studies on the mechanism of the antiviral activity of ribavirin against reovirus. Virology 168:147–158.

44. Borsa J, Long DG, Copps TP, Sargent MD, Chapman JD. 1974. Reovirus transcriptase activation *in vitro*: further studies on the facilitation phenomenon. Intervirology 3:15–35.

45. Borsa J, Long DG, Sargent MD, Copps TP, Chapman JD. 1974. Reovirus transcriptase activation *in vitro*: involvement of an endogenous uncoating activity in the second stage of the process. Intervirology 4:171–188.

46. Chandran K, Farsetta DL, Nibert ML. 2002. Strategy for nonenveloped virus entry: a hydrophobic conformer of the reovirus membrane penetration protein m1 mediates membrane disruption. Journal of Virology 76:9920–9933.

47. Farsetta DL, Chandran K, Nibert ML. 2000. Transcriptional activities of reovirus RNA polymerase in recoated cores. Initiation and elongation are regulated by separate mechanisms. J Biol Chem 275:39693–701.

48. Banerjee AK, Shatkin AJ. 1970. Transcription in vitro by reovirus-associated ribonucleic acid-dependent polymerase. Journal of Virology 6:1–11.

49. Dryden KA, Wang G, Yeager M, Nibert ML, Coombs KM, Furlong DB, Fields BN, Baker TS. 1993. Early steps in reovirus infection are associated with dramatic changes in supramolecular structure and protein conformation: analysis of virions and subviral particles by cryoelectron microscopy and image reconstruction. Journal of Cell Biology 122:1023–1041.

50. Reinisch KM, Nibert ML, Harrison SC. 2000. Structure of the reovirus core at 3.6 Å resolution. Nature 404:960–967.

51. Tenorio R, Fernandez de Castro I, Knowlton JJ, Zamora PF, Lee CH, Mainou BA, Dermody TS, Risco C. 2018. Reovirus sigmaNS and muNS Proteins Remodel the Endoplasmic Reticulum to Build Replication Neo-Organelles. MBio 9.

52. Furuichi Y, Muthukrishnan S, Shatkin AJ. 1975. 5′-Terminal M7G(5′)ppp(5′)Gmp in vivo: identification in reovirus genome RNA. Proceedings of the National Academy of Sciences USA 72:742–745.

53. Silverstein SC, Astell C, Levin DH, Schonberg M, Acs G. 1972. The mechanism of reovirus uncoating and gene activation *in vivo*. Virology 47:797–806.

